# HAPLN1 is a driver for peritoneal carcinomatosis in pancreatic cancer

**DOI:** 10.1101/2022.05.30.493185

**Authors:** Lena Wiedmann, Francesca De Angelis Rigotti, Nuria Vaquero-Siguero, Elisa Donato, Elisa Espinet, Iris Moll, Elisenda Alsina-Sanchis, Ronja Mülfarth, Margherita Vacca, Jennifer Gerwing, Andreas Trumpp, Andreas Fischer, Juan Rodriguez-Vita

**Author notes:** Equal contribution (Lena Wiedmann, Francesca De Angelis Rigotti, Andreas Fischer, Juan Rodriguez-Vita).

## Abstract

Pancreatic ductal adenocarcinoma (PDAC) frequently metastasizes into the peritoneum, which contributes to poor prognosis. Metastatic spreading is promoted by cancer cell plasticity, yet its regulation by the microenvironment is incompletely understood. Here we show that the presence of hyaluronan and proteoglycan link protein-1 (HAPLN1) in the extracellular matrix enhances tumor cell plasticity and PDAC metastasis. Bioinformatic analysis showed that HAPLN1 expression is enriched in the basal PDAC subtype and associated with worse overall patient survival. In a mouse model for peritoneal carcinomatosis, HAPLN1-induced immunomodulation favored a more permissive microenvironment, which accelerated the peritoneal spread of tumor cells. Mechanistically, HAPLN1, via hyaluronic acid synthesis and signaling, promoted adoption of a highly plastic cancer cell state, facilitating EMT, stemness, invasion and immunomodulation in a paracrine manner. Extracellular HAPLN1 modifies cancer cells as well as fibroblasts, rendering them immunomodulatory. We identify HAPLN1 as a prognostic marker and a driver for peritoneal metastasis in PDAC.

## Introduction

Pancreatic ductal adenocarcinoma (PDAC) ranks among the most lethal cancer entities, with late diagnosis and early onset of metastatic spread as key contributors to its poor survival rate (1). PDAC metastasis most commonly occurs in the liver (2); however, many patients suffer from peritoneal carcinomatosis, which is a major, but understudied cause of morbidity and mortality of patients with no effective treatment options (3). Indeed, at time of diagnosis around 9 % of PDAC patients present with peritoneal metastasis, while the rate at death raises up to 25-50 % (3). Therefore, understanding the mechanisms of peritoneal dissemination is of critical clinical need to shed a light into new treatment options. Metastatic tumor cells need to acquire certain features, which allow them to survive and grow out within a distant hostile microenvironment and to escape the immune response. All these characteristics can be controlled by unlocking cellular plasticity, an event recently incorporated to the hallmarks of cancer (4). Cellular plasticity is characterized by epithelial-mesenchymal plasticity (EMP) and features of cancer cell stemness (5). Plasticity not only makes cancer cells more prone for invasion and adaption to the microenvironment, but also protects them from apoptosis, immune attack and chemotherapy (6).

Despite the cancer cell intrinsic features, a key regulator of metastasis is the microenvironment that tumor cells are facing during their journey to the metastatic site (7). The tumor microenvironment (TME) consists of several cellular and non-cellular components, including cancer-associated fibroblasts (CAFs), endothelial cells, immune cells and extracellular matrix (ECM). In PDAC, the TME is strongly desmoplastic, with substantial accumulation of ECM components. Hyaluronic Acid (HA), one of the major components of the ECM, facilitates tumor progression and metastasis through promoting EMP, invasion, immunomodulation and therapy resistance (8). CAFs are the main producers of ECM, however their additional role as immunomodulators has been increasingly recognized. By the expression of different cytokines, growth factors, and immunomodulatory molecules, CAFs impact on the recruitment, differentiation and polarization of innate and adaptive immunity (9).

When metastasizing into the peritoneum, disseminated tumor cells predominantly colonize milky spots of the omental fat pad, a metastatic niche with high nutrient levels and immune cells with more resolving, antiinflammatory features (mostly resident macrophages and B cells) (10). For instance, omental resident macrophages were proven crucial for metastatic progression and immunomodulation (11).

From the omentum, cancer cells colonize the whole peritoneum. Suspended in the peritoneal cavity, tumor cells face different challenges, like anoikis, which can be prevented by cell cluster (spheroid) formation, maintenance of mesenchymal and stemness state (e.g. via STAT3 signaling) and/or survival signaling like PI3K, MEK–ERK, or NF-κB–FAK (3). Additionally, tumor cells are facing tumoricidal immune cells. However, tumor-associated macrophages (TAMs), which are the predominant immune population in the diseased peritoneum, allow peritoneal metastasis from multiple tumor entities (11,12). The phenotypic switch of resident tumoricidal peritoneal macrophages towards the TAM phenotype is affected by factors such as tumor-derived HA (13). Thus, targeting HA is not only promising for localized, but also metastatic PDAC. However, clinical trials targeting HA with the hyaluronidase PEGPH20 failed in stage-3 (14), therefore new treatment options are of utmost need.

Hyaluronan and proteoglycan link protein-1 (HAPLN1) is a HA and chondroitin sulfate proteoglycan (CSPG) crosslinker in the ECM, with so far poorly understood roles in cancer. Recently, CAF-derived HAPLN1 was found to fuel tumor cell invasion in gastric cancer and was associated with worse overall survival in pleural mesothelioma and drug resistance in multiple myeloma (15–17). In contrast, HAPLN1 expression was associated with reduced disease progression in melanoma and colorectal cancer (18–20). Additionally, in hepatocellular carcinoma HAPLN1 is expressed by tumor cells and associated with epithelial-to-mesenchymal transition (EMT) (21). Nevertheless, its functional role in PDAC remains elusive

In this study we identified HAPLN1 as one of the most upregulated genes in PDAC compared to adjacent tissue. We identify HAPLN1 as a novel mediator of peritoneal dissemination in PDAC, by inducing a highly plastic phenotype in cancer cells, which leads to a pro-tumoral metastatic niche.

## Results

### HAPLN1 is upregulated in PDAC tissue and associated with worse disease outcome

To unravel novel mediators of HA-mediated PDAC progression, we analyzed publicly available data sets of PDAC and adjacent tissue from patients (Cao et al. 2021, GSE62452, (22,23)). We performed gene set enrichment analysis (GSEA) using a gene set for HA-binding (Gene Ontology for “Hyaluronic Acid Binding”) to compare its expression between tumorous and adjacent tissue when both data were available (**Figure 1A, Suppl. Fig. 1A**). Interestingly, *HAPLN1* was the most enriched gene in one of the analyses, and within the leading edge in both data sets.

**Figure 1:**
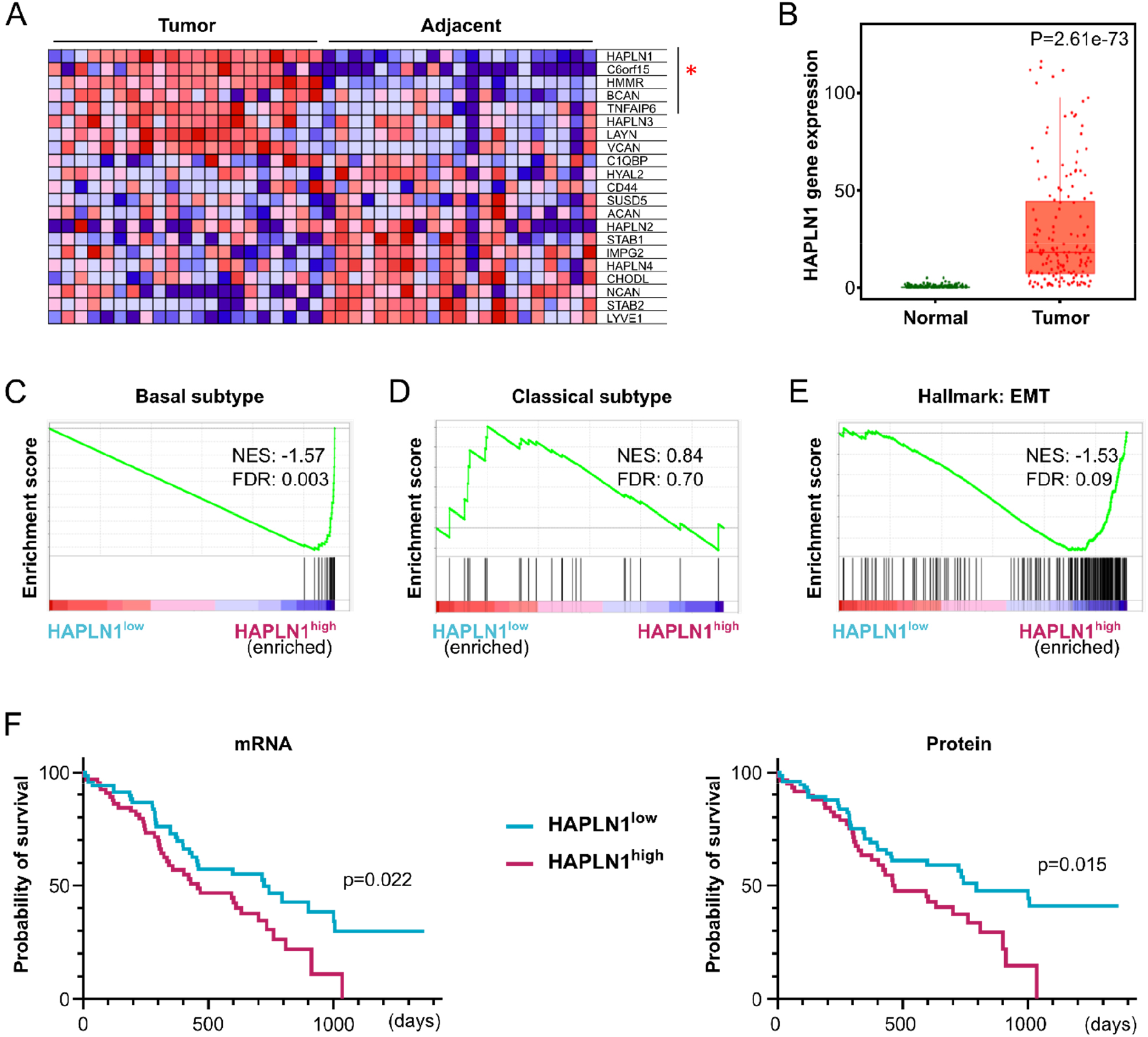
HAPLN1 is upregulated in PDAC tissue and associated with worse disease outcome. **A**. GSEA of data set published in Cao et al. (2021) (23) (Matched tumor and adjacent tissue samples. n=21) on “Gene ontology for Hyaluronic Acid binding”. Leading edge genes significantly deregulated are marked with red star. **B.** HAPLN1 expression levels in normal adjacent or PDAC tumor tissue according to TNMplot. **C-E.** GSEA on data set of Cao et al. (2021). Before analysis PDAC patients were divided in HAPLN1 high/low according to mean HAPLN1 expression. Gene sets of “Basal subtype” (**C**), “Classical subtype” (**D**) or “Hallmark of Epithelial-to-Mesenchymal transition” (**E**). n=140. **F**. Overall survival (OS) of PDAC patients stratified in two cohorts according to their mean HAPLN1 mRNA or protein expression levels. n=135. For OS log-rank (Mantel-Cox) test was applied.

The significant upregulation of *HAPLN1* gene expression in tumorous tissue compared to healthy was confirmed using the TNMplot database (24) (**Figure 1B**).

Further, we classified the tumorous tissue samples as *HAPLN1^high^* or *HAPLN1^low^* based on mean *HAPLN1* expression level. By GSEA, we discovered a significant enrichment of the gene set for the basal PDAC subtype in the *HAPLN1*^high^ samples, while the gene set of classical PDAC subtype was enriched in *HAPLN1^low^* ones (**Figure 1C,D; Suppl. Figure 1B,C**). The basal subtype is less differentiated and patients have a lower overall survival than those with the classical subtype (25).

Moreover, EMT-related gene sets were significantly enriched in *HAPLN1*^high^ as observed by GSEA of the gene set “Hallmark: Epithelial-to-mesenchymal transition” (**Figure 1E, Suppl. Figure 1D**), reinforcing our hypothesis of HAPLN1 as a novel mediator of PDAC progression.

Indeed, when addressing if HAPLN1 expression is associated with disease outcome, we found that overall survival of patients was significantly reduced in the *HAPLN1^high^* group (**Figure 1F**). Moreover we confirmed this association in patients with higher HAPLN1 protein content in tumorous tissue (**Figure 1F**).

### HAPLN1 induces EMT and ECM remodeling in cultured PDAC cells

To understand the possible functional role of HAPLN1 in PDAC, we overexpressed HAPLN1 in a murine PDAC tumor cell line derived from KPC (Kras^G12D^; Trp53^R172H^; Elas-CreER (26)) mice. KPC cells have very low HAPLN1 expression under normal cell culture conditions (**Suppl. Fig. 2A**). Forced expression of HAPLN1 did not change cell proliferation (**Suppl. Fig. 2B**). However, KPC-HAPLN1 cells had a more mesenchymal appearance compared to KPC cells that grew in islet-like cell associations (**Suppl. Figure 2C**). Since HAPLN1 acts as crosslinker between HA and proteoglycans in the ECM, we addressed if KPC-HAPLN1 cells showed changes in their HA production. Interestingly, significantly more HA was detected in the supernatant of KPC-HAPLN1 cells (**Figure 2A**). This was caused by an upregulation of HA-Synthase 2 (HAS2), while the other synthases (*Has1, Has3*) remained unchanged (**Figure 2B,C**). HAS2 is the main producer of high molecular weight (HMW)-HA and a known inducer of EMT (27). Thus, we analyzed EMT traits by investigating changes of mRNA and protein levels of mesenchymal and epithelial markers. This confirmed a more mesenchymal state of KPC-HAPLN1 cells, with mesenchymal markers like TWIST1 and LRRC15 being upregulated, while the epithelial marker E-Cadherin was downregulated (**Figure 2D,E**). Moreover, increased motility, a feature of a more mesenchymal phenotype, was observed (**Suppl. Fig. 2D, Suppl. Video 1,2)**.

**Figure 2:**
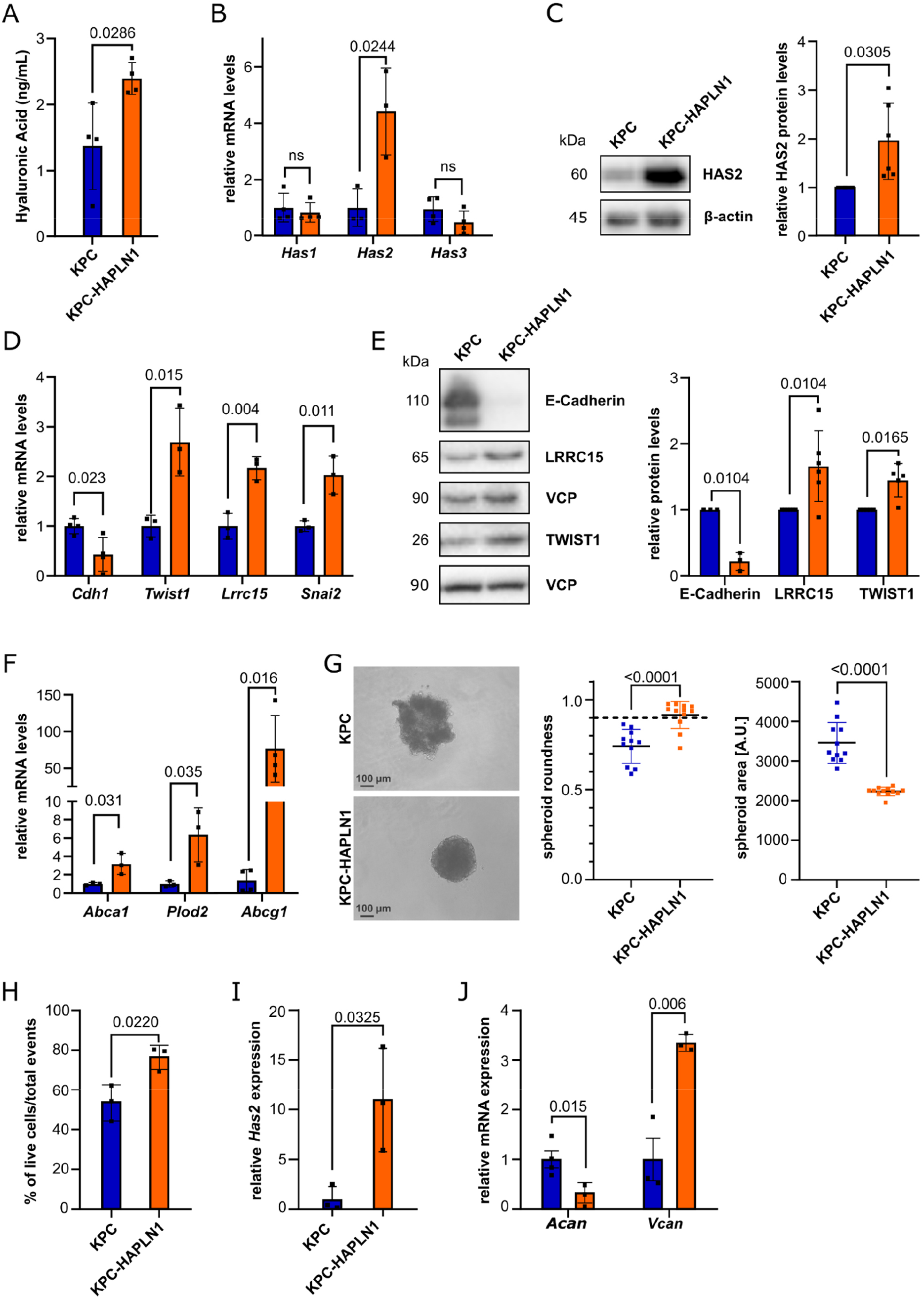
HAPLN1 induces EMT and ECM remodeling in vitro. **A**. ELISA-like assay for Hyaluronan (HA) in the serum-free supernatant of KPC and KPC-HAPLN1 cells. n=4. **B**. qRT-PCR analysis of HA-Synthase (Has) expression levels. n=3-4. **C**. Confirmation of increased HAS2 levels by Western blot analysis. Quantification by normalization to house keeper β-Actin. n=6. **D**. mRNA expression of epithelial and mesenchymal markers comparing KPC and KPC-HAPLN1 cells. n=3-4. **E**. Western blot analysis of epithelial and mesenchymal markers on protein lysates. Quantification by normalization to house keeper VCP. n=3-6. **F**. mRNA expression of stemness markers by qRT-PCR. n=3-4. **G**. Cells seeded in ultra-low attachment plates. Pictures taken after 48 h. Quantification of area and roundness as measures for proper spheroid formation. Scale bar: 100 μm. n=11-12. **H**. Flow cytometric analysis of DAPI^-^ events using spheroids formed for 48 h. n=3. **I.+J.** Gene expression analysis of Has2 (**I.**) and HAPLN1-associated proteoglycans (**J.**) in spheroids. n=3-4. Plots are shown as mean±SD. Data points represent independent biological replicates. For panels A, B, D, F-J unpaired two-tailed T test was applied, for panels C+E paired two-tailed T test.

Since HA does not only affect EMT, but also stemness (28), we investigated the effect of HAPLN1 on the expression of stemness-related genes *Abca1, Plod2* and *Abcg1* (**Figure 2F**). The significant upregulation of all these markers in KPC-HAPLN1 cells prompted us to assess spheroid formation as a readout for stemness. KPC-HAPLN1 cells were capable of forming proper and round spheroids in 3D culture, while KPC cells only formed aggregate-like structures (**Figure 2G**). Measuring the area and roundness as indicators of complete spheroids, we could confirm the optimal spheroid formation capacity of KPC-HAPLN1 cells as shown by a significant decrease of their area and a roundness of >0.9 (**Figure 2G**). Matching with the previous results, spheroids formed by KPC-HAPLN1 cells contained significantly more alive cells than spheroids formed by KPC cells, indicating an improved 3D organization inside the spheroid (**Figure 2H**).

The linker function of HAPLN1, crosslinking HA to proteoglycans (**Suppl. Figure 2E**), suggests that the improved spheroid formation could be mediated by changes in their ECM composition. Thus, we addressed gene expression of the HA synthase *Has2*, as well as of the proteoglycans aggrecan (*Acan*) and versican (*Vcan*) in spheroids. *Has2* and *Vcan* levels were significantly upregulated, while *Acan* levels were strongly reduced (**Figure 2I,J**). In summary, these data demonstrate that HAPLN1 expression remodels the ECM and promotes EMT and stemness of KPC cells.

### HAPLN1 promotes invasion

EMP, motility, and stemness are signs of cellular plasticity. Another feature of high plasticity is invasion. To evaluate the role of HAPLN1 expression in invasion, we embedded tumor cell spheroids into Matrigel and evaluated their invasive potential (**Figure 3A**). After the addition of Matrigel, also KPC cells formed round spheroids, confirming the important role of ECM in this process (**Figure 3B**). However, these cells were not able to invade the Matrigel. In contrast, KPC-HAPLN1 cells robustly invaded the Matrigel (**Figure 3B**).

**Figure 3:**
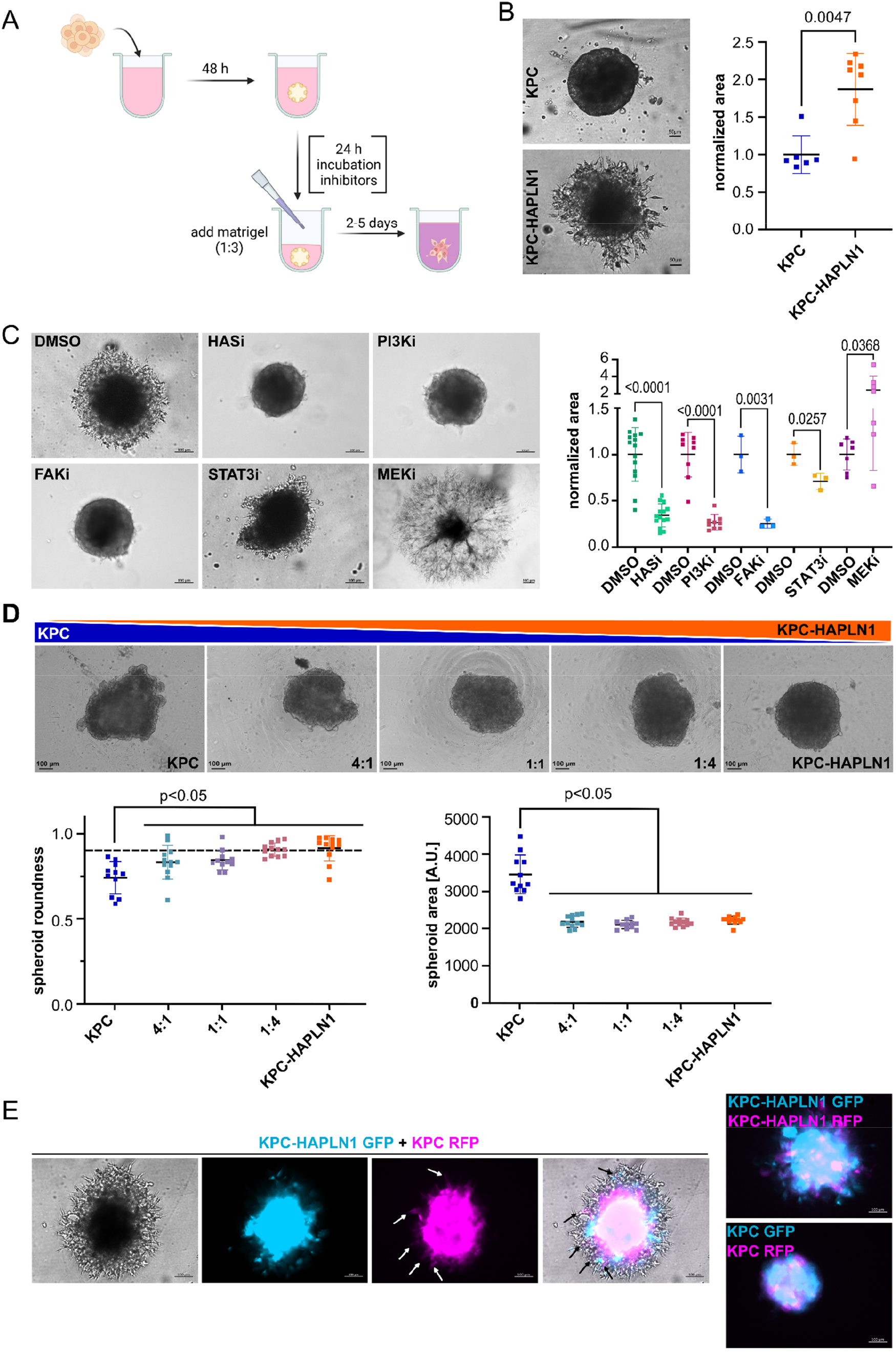
HAPLN1 fuels invasion and acts in a paracrine manner. **A**. Schematic overview of spheroid formation, possible treatment and embedding into Matrigel to assess invasive capacity. **B**. Invasion of KPC or KPC-HAPLN1 cells 48 h after embedding of the spheroids into Matrigel. The occupied area of spheroids embedded into Matrigel is quantified as a readout of invasion (n=6-8) and representative images are shown. Scale bar: 50 μm. **C**. KPC-HAPLN1 spheroids were pre-treated with inhibitors before embedding into Matrigel. Invasion was assessed 72-96 h later. HASi=500 μM 4-Methylumbelliferrone; PI3Ki=10 μM LY294002; FAKi=10 μM GSK2256098; STAT3i=25 μM Stattic; MEKi=50 μM PD98059. Representative images (scale bar: 100 μm) and quantification by assessing occupied area (n=3-14) are shown. **D**. KPC and KPC-HAPLN1 cells were mixed at different ratios. Images are representative of each condition. Scale bar: 100 μm. Spheroid roundness and area are quantified by Fiji software. N=11-12. **E**. KPC and KPC-HAPLN1 cells were labeled with GFP or RFP via lentiviral vectors. Invasion was assessed after 48-72 h. Scale bar: 100 μm. Graphs represent mean±SD. Each data point represents a biological replicate. Unpaired two-tailed T test was used for analysis.

HA has been shown to act through CD44 activation (28). In order to investigate the role of this interaction during HAPLN1-induced invasion we inhibited several components of the HA/CD44 signaling pathway. We inhibited HA synthase activity as well as PI3K, FAK, STAT3 and MEK which are transducers of CD44 signaling (28), in KPC-HAPLN1 spheroids. Invasion was dependent on HA production, as well as PI3K, FAK and STAT3 signaling, while MEK inhibition even increased the invasive potential of cells (**Figure 3C**).

Interestingly, also the treatment of KPC cells with the MEK inhibitor increased their invasion capacity, corroborating that MEK activation in this setting negatively regulates invasion (**Suppl. Figure 3A**). In keeping with this, we found that KPC-HAPLN1 cells had FAK and STAT3 signaling activated as shown by their increased phosphorylation (**Suppl. Figure 3B**). This recapitulated previously published studies showing that HA increases the invasive capabilities of cells, through PI3K, FAK and STAT3 signaling (28) and that MEK inhibition induces FAK and STAT3 signaling, proving again their essential role in HAPLN1-induced invasion (29).

### HAPLN1 acts in a paracrine manner

Since HAPLN1, as well as the affected ECM components HA, VCAN and ACAN are extracellular, we evaluated the potential paracrine effects of HAPLN1 expression. For this, we mixed KPC and KPC-HAPLN1 cells at different ratios and assessed spheroid formation capacity (**Figure 3D**). Here, already in a 4:1 (KPC:KPC-HAPLN1) ratio, spheroid formation capacity was significantly improved, measured by area and roundness (**Figure 3D**).

To understand whether these paracrine effects of HAPLN1 could also impact on invasion, we labelled KPC and KPC-HAPLN1 with RFP and GFP respectively and co-cultured them as spheroids and again embedded them into Matrigel. Interestingly, KPC cells invaded this Matrigel only when co-cultured with KPC-HAPLN1 cells (**Figure 3E**). Additionally, we used adenoviral overexpression of GFP or mCherry to exclude an effect of lentiviral expression in the obtained results (**Suppl. Figure 3C**). In conclusion, our data demonstrate that the presence of HAPLN1 in the ECM is sufficient to increase tumor cell malignancy.

### HAPLN1 induces tumor cell high-plasticity *in vivo*

To explore the effects of HAPLN1 on EMT, stemness and invasion *in vivo*, we orthotopically implanted KPC or KPC-HAPLN1 cells into wildtype C57Bl6/J mice. Even though we injected only 10,000 cells, tumor size reached the regulatory termination criteria before signs of invasion and metastasis could be observed (data not shown). For this reason, we decided to use a model for peritoneal carcinomatosis, injecting RFP and luciferase-expressing KPC and KPC-HAPLN1 cells intraperitoneally (i.p.), to mimic an advanced tumor stage. This model is frequently used to study peritoneal dissemination and metastasis in other abdominal cancers, such as ovarian and gastric cancer, where tumor cells settle in the omental fat pads before spreading throughout the peritoneal cavity (11).

When analyzing the cell composition of the tumor masses formed in the omentum by flow cytometry and immunofluorescence, we detected that immune cell distribution within the tissue was very heterogeneous (**Suppl. Figure 4A,B**), with immune deserted tumor centers and high to moderate presence on the borders (**Suppl. Figure 4B**), which prevented further quantification. Therefore, we next analyzed mRNA expression levels of bulk tumor samples to get a deeper understanding of the milieu. In KPC-HAPLN1 tumors, there was higher expression of the immunomodulatory markers *Cd274* (PD-L1) and MHC-II complex component *H2-Ab1* compared to KPC control tumors (**Suppl. Figure 4C**).

On the other hand, flow cytometry showed an increase in the presence of PDPN^+^ fibroblasts (PDPN^+^/CD3T^-^/CD45^-^/RFP^-^) in KPC-HAPLN1 tumor-bearing mice (**Suppl. Figure 4A**). To understand the localization of these fibroblasts, we performed immunofluorescence analysis. When staining for αSMA as an activated fibroblast marker, we confirmed the increase of fibroblasts in KPC-HAPLN1 tumors (**Figure 4A**). As observed with flow cytometry, endothelial cell (CD31^+^) presence was unchanged between both groups (**Figure 4A**).

**Figure 4:**
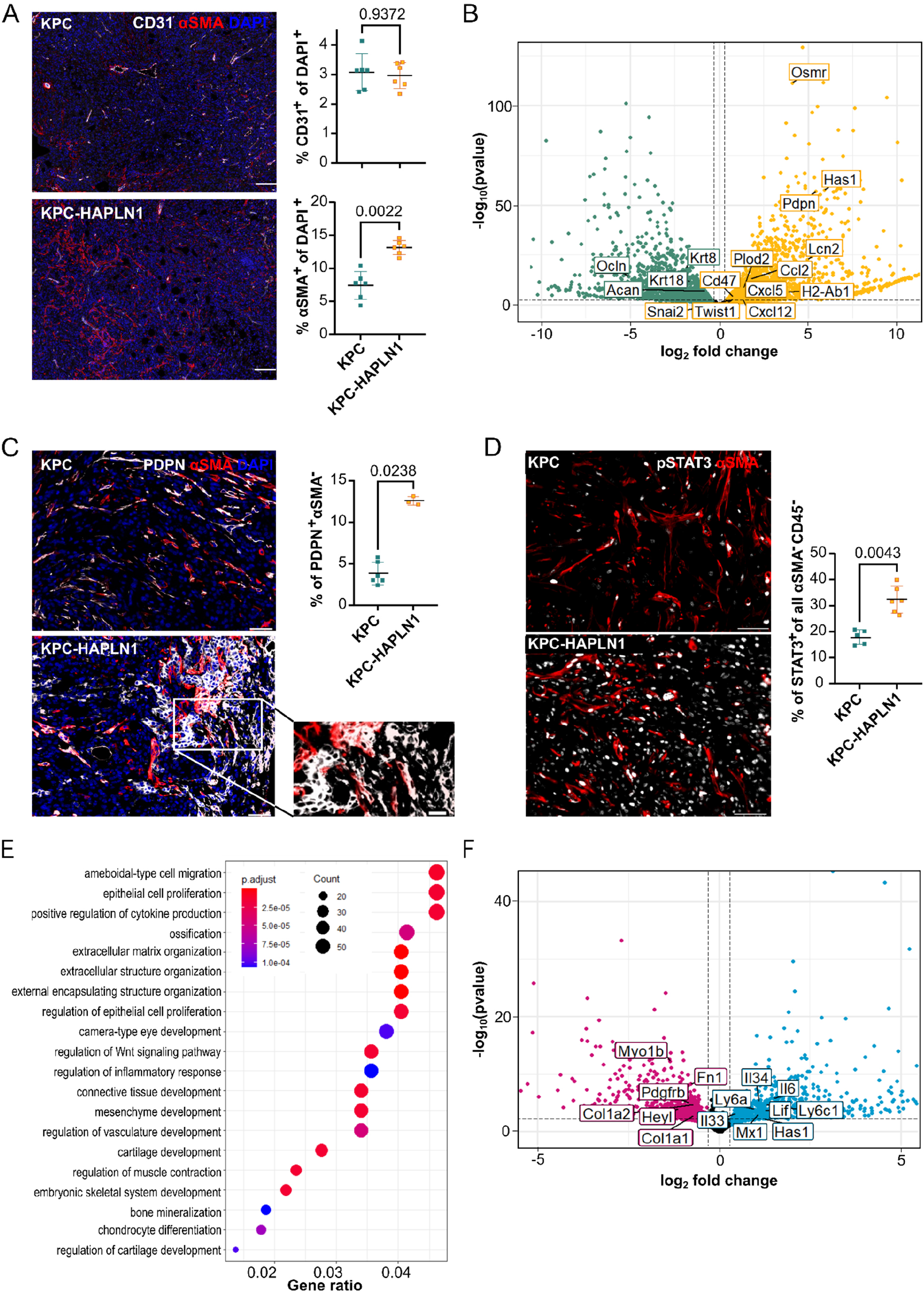
HAPLN1 induces tumor cell high-plasticity in vivo. Mice were injected with KPC or KPC-HAPLN1 cells intraperitoneally to mimic peritoneal metastasis. Solid tumors formed in the omental region. **A**. CD31/αSMA immunofluorescence staining on tumors. Representative images and quantification of CD31^+^ cells and αSMA^+^ cells are shown. Scale bar: 50 μm. n=6. **B**. Volcano plot obtained from RNA sequencing on tumor cells that were isolated from KPC or KPC-HAPLN1 solid tumors. Each dot represents one gene, with green dots upregulated in KPC and yellow dots upregulated in KPC-HAPLN1 cells. Some genes of interest were labeled. n=3. **C**. Podoplanin (PDPN)/αSMA staining on tumors. Representative images and quantification of PDPN^+^ αSMA^-^ cells are shown. Scale bar: 50 μm, zoom: 20 μm. n=3-6. **D**. pSTAT3/αSMA staining on tumors. Representative images and quantification of STAT3^+^ tumor cells are shown. n=5-6. **E**. Gene Ontology for ‘Biological Process’ of genes significantly upregulated in KPC-HAPLN1 cells compared to KPC. n=3. **F**. RNA sequencing of PDPN^+^ isolated cancer-associated fibroblasts from KPC or KPC-HAPLN1 tumors. Pink dots mark genes significantly up in CAFs from KPC, blue dots genes significantly up in CAFs from KPC-HAPLN1 tumors. Some genes of interest are labeled. n=3. Graphs represent mean±SD. Data points indicate independent biological replicates. For image analysis non-parametric Mann-Whitney U test was used.

To assess EMT-related processes even in the presence of higher fibroblast numbers, we sorted tumor cells (DAPI^-^/CD45^-^/CD31^-^/RFP^+^) by flow cytometry and investigated their expression profile by RNA sequencing. Principal component analysis (PCA) confirmed clustering of the two different groups, with almost all of the variation attributable to HAPLN1 expression (**Suppl. Figure 4D**). When extracting significantly deregulated genes between KPC and KPC-HAPLN1 we found several markers of EMT (e.g. *Snai2*, *Twist1*, *Pdpn*, *Ocln*, *Krt8*, *Krt18*), stemness (e.g. *Cxcl12*, *Plod2*, *Osmr*), ECM remodeling (*Has1*, *Acan*) and immunomodulation (e.g. *H2-Ab1* (MHC-II), *Ccl2*, *Lcn2*, *Cxcl5*, *Cd47*) deregulated in KPC-HAPLN1 cells (**Figure 4B**). We validated the upregulation of PDPN and MHC-II by flow cytometry. The majority of KPC-HAPLN1 cells expressed PDPN and MHC-II on their plasma membrane compared to the lack of staining on nearly all KPC cells (**Suppl. Fig. 4E**). The upregulation of PDPN in tumor cells was additionally confirmed by immunofluorescence staining (**Figure 4C**).

GSEA comparing KPC and KPC-HAPLN1 transcription profiles revealed that gene sets for “Hallmark: IL6-JAK-STAT3 signaling” and “Canonical Wnt signaling pathway” were significantly enriched in the KPC-HAPLN1 cells. IL6-JAK-STAT3 signaling was already shown to promote metastatic progression, including peritoneal metastasis (30,31), while the enrichment of Wnt signaling corroborates a more stemness-promoting milieu (32) (**Suppl. Fig. 4F**). Immunofluorescence staining of pSTAT3 confirmed the increase in STAT3 activation in tumor cells (αSMA^-^ /CD45^-^) from KPC-HAPLN1 tumor-bearing mice (**Figure 4D**).

Finally, Gene Ontology (GO) analysis confirmed our previous results with terms like “ameboidal-type cell migration”, “positive regulation of cytokine production”, “extracellular matrix organization”, “regulation of Wnt signaling pathway” and “regulation of inflammatory response” enriched in KPC-HAPLN1 cells (**Figure 4E**). Given the strong presence of CAFs in these tumors, we decided to investigate also their expression profile by RNAseq. We sorted fibroblasts (DAPI^-^/CD45^-^/CD31^-^ /RFP^-^/PDPN^+^) from KPC and KPC-HAPLN1 tumors by flow cytometry. Interestingly, the expression profile of CAFs clustered in the same way as the tumors from which they were obtained (**Suppl. Figure 4G**), indicating that HAPLN1 presence in the ECM is sufficient to deploy a distinguishable phenotype also in stromal cells. We found that fibroblasts of KPC-HAPLN1 tumors significantly upregulated immunomodulatory and inflammation-related genes like *Lif*, *Il6* and others (e.g. *Il33*, *Il34*, *Ly6a*, *Ly6c1*, *Mx1*, *Has1*) compared to fibroblasts in KPC tumors. The increased expression of *Il6* suggests that fibroblasts could contribute to the increased STAT3 phosphorylation observed in KPC-HAPLN1. In contrast, fibroblasts from KPC tumors had the expression of ECM components like *Col1a1*, *Col1a2*, *Fn1* and others (e.g. *Pdgfrb*, *Myo1b*, *Heyl*) significantly increased (**Figure 4F**).

Taken together our data show that the presence of HAPLN1 alters CAF biology in a paracrine manner, in addition to promoting a highly plastic tumor cell state.

### HAPLN1 modifies the immune microenvironment

Since both, tumor cells and CAFs from KPC-HAPLN1 tumors showed changes in markers of inflammation and immunomodulation, we analyzed the blood from tumor-bearing mice to evaluate whether the inflammatory milieu could even be detected systemically. We noticed a significant induction of IFNγ and inhibin-A (INHBA) protein levels in serum of KPC-bearing mice compared to control, which were abolished nearly to baseline in KPC-HAPLN1 tumor-bearing mice (**Figure 5A**). Moreover, the immune checkpoint molecule PD-L1 was upregulated in KPC-HAPLN1 compared to control KPC-injected mice, although this difference did not reach statistical significance (**Figure 5A**). In addition, the type-2 inflammatory mediator IL33 was only upregulated in KPC-HAPLN1 tumor bearing mice (**Figure 5A**). These data support the hypothesis that HAPLN1 promotes an anti-inflammatory state. To understand the impact of these systemic changes on the peritoneal cavity, we analyzed the peritoneal lavage (PL) of tumor-bearing mice. Firstly, we detected a strong increase in the number of cells present in the peritoneal cavity of KPC-HAPLN1 compared to control KPC-bearing mice (**Figure 5B**). When analyzing the immune compartment, we found a much higher percentage of macrophages and B lymphocytes in the PL of KPC-HAPLN1 compared to KPC tumor-bearing mice, while the percentages of monocytes and neutrophils were significantly lower (**Figure 5C, Suppl. Figure 5A**). We also observed a drastic reduction in eosinophil percentages, while helper, effector and regulatory T cell populations were unchanged (**Suppl. Figure 5A,B**). This is a typical feature of peritoneal metastasis, where myeloid cells are the main immune component and are in charge of immunosuppression (10).

**Figure 5:**
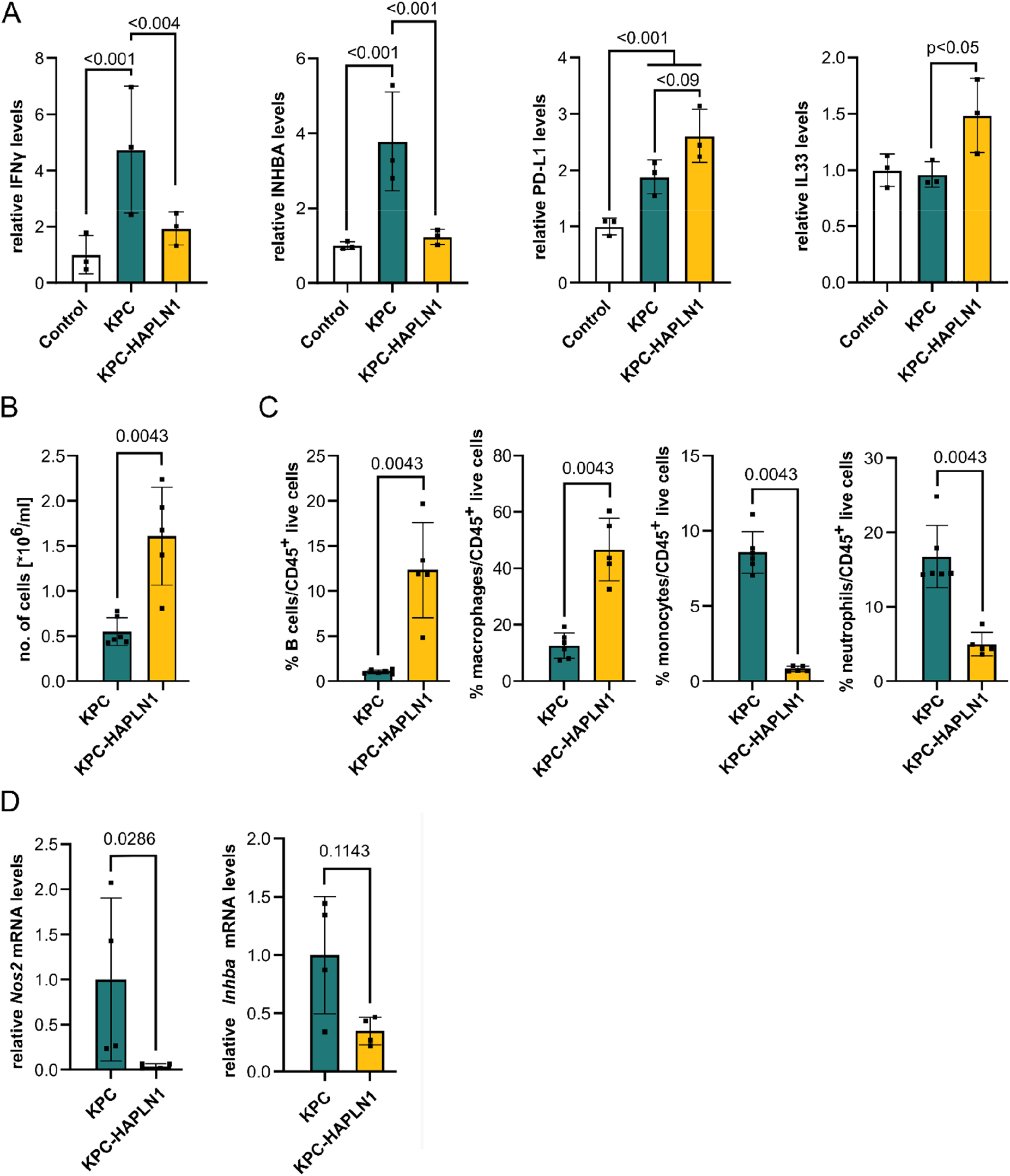
HAPLN1 modifies the immune microenvironment. **A**. Serum of healthy, or KPC/KPC-HAPLN1-injected mice at day 11 of tumor growth was analyzed using the scioCD protein array. Relative protein levels displayed. n=3. **B**. Number of cells per milliliter in the peritoneal lavage of mice after 11 days of tumor growth. n=5-6. **C**. Immune cell populations in the peritoneal lavage assessed by flow cytometry. n=5-6. **D**. Macrophages were isolated from the peritoneum by cell sorting. qRT-PCR analysis of inflammatory marker genes. n=4. Graphs represent mean±SD. Data points indicate independent biological replicates. Non-parametric Mann-Whitney U test was used.

Tumor-associated macrophages (TAMs) are one of the most important mediators of peritoneal immunity and their proportions are directly associated with worse progression in different cancer entities (33,34). To understand if TAMs in KPC-HAPLN1 tumor-bearing mice could be contributing to tumor progression as well, we isolated them and performed qRT-PCR analysis. This revealed reduced expression levels of pro-inflammatory markers like *Nos2* (iNOS) or *Inhba* in macrophages derived from KPC-HAPLN1 mice (**Figure 5D**). In addition, when stimulating bone marrow-derived macrophages (BMDMs) with cancer cell conditioned medium, we detected significantly higher levels of the anti-inflammatory macrophage marker *Arg1* (**Suppl. Figure 5C**) when KPC cells expressed HAPLN1, indicating that HAPLN1 promotes TAM education in a paracrine manner.

### HAPLN1 facilitates peritoneal colonization by tumor cells

The data presented so far indicate that HAPLN1 expression in PDAC cells induces a more aggressive, highly plastic state and a more tumor-permissive niche within the peritoneal cavity. For this reason, we decided to further investigate peritoneal dissemination of tumor cells.

We detected increased luminescence in PL from KPC-HAPLN1 compared to that from KPC tumor-bearing mice, which directly correlates with increased presence of tumor cells in peritoneal fluid (**Figure 6A**). The increased luminescence in the PL was however not attributable to an increased size of the tumor masses present in the omentum (**Suppl. Figure 6A**). Flow cytometry of PL confirmed that there were more detached RFP^+^ tumor cells in KPC-HAPLN1 compared to control KPC tumor-bearing mice (**Figure 6B**).

**Figure 6:**
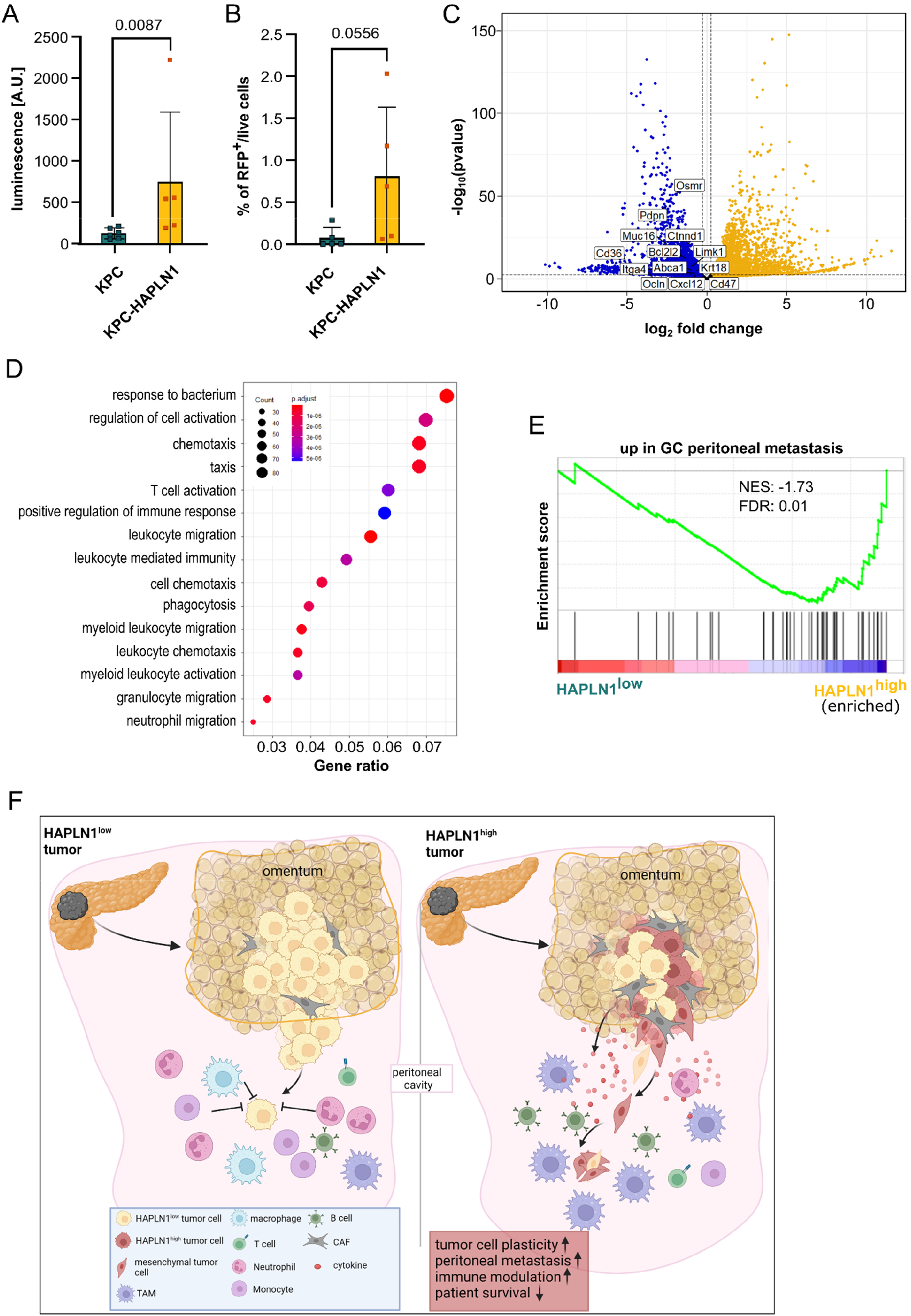
HAPLN1 facilitates peritoneal colonization by tumor cells. **A**.Ex vivo luminescence measurement on peritoneal lavage of KPC and KPC-HAPLN1 mice 11 days after tumor injection. n=5-6. **B**. RFP^+^ tumor cells detected by flow cytometry in the peritoneal lavage of mice at day 11. **C**. RNA sequencing results comparing KPC-HAPLN1 cells from solid tumor or in suspension in peritoneal lavage. Blue dots mark genes upregulated in cells in suspension, yellow dots genes upregulated in cells in tumor mass. Some genes of interest are highlighted. n=3. **D**. Gene Ontology of ‘Biological Process’ on genes upregulated in KPC-HAPLN1 cells which detached into the peritoneal cavity. **E**. GSEA on the PDAC patient data set of Cao et al (2021), divided in high or low HAPLN1 expression by the mean HAPLN1 expression. A gene set for genes upregulated in gastric cancer (GC) peritoneal metastasis was used. n=140. **F**. Graphical abstract of the study presented. Graphs represent mean±SD. Data points indicate independent biological replicates. Non-parametric Mann-Whitney U test was used for panel A+B.

One can expect that such detached cancer cells contribute to tumor spread within the whole peritoneal cavity. As such, we aimed at deciphering how these cells differ from the cancer cells within solid tumor masses. We isolated KPC-HAPLN1 tumor cells from the PL by cell sorting (DAPI^-^CD45^-^/RFP^+^) and performed RNA sequencing. Cells from solid tumor masses or in a detached state separated clearly when performing principal component analysis (PCA) (**Suppl. Figure 6B**). Among the genes expressed significantly higher in the detached population, we found stemness-associated genes like *Cxcl12* and *Abca1*, apoptosis-inhibitors like *Birc2* and *Bcl2l1*, peritoneal metastasis-associated genes, e.g. *Ctnnd1, Itga4, Limk1*, and immunomodulatory genes like *Pdpn, Cd47, Cd36, Osmr* and *Muc16*. Additionally, the epithelial cell markers *Krt18* and *Ocln* were highly expressed in detached vs. solid tumor cells. This suggests the acquisition of a more epithelial state, a process which was shown to be important for the attachment to and colonization of secondary organs (**Figure 6C**, (35)). As expected, when performing GO term analysis, almost all terms enriched in disseminated tumor cells were associated with immunity, inflammation and migration (**Figure 6D**), suggesting that disseminated KPC-HAPLN1 cells could have cell-intrinsic features to modulate immune response and shape the microenvironment they are facing.

In order to validate these results, we analyzed the abovementioned patient datasets classified as *HAPLN1^high^* and *HAPLN1^low^* and performed GSEA using a gene set of differentially expressed genes in peritoneal metastasis in patients with gastric cancer

(36). In both data sets, the peritoneal metastasis gene set was enriched in the *HAPLN1*^high^ patient cohort, reinforcing the hypothesis that HAPLN1 increases the potential of tumors to develop peritoneal carcinomatosis (**Figure 6E, Suppl. Figure 6C**).

Overall, we conclude that HAPLN1 expression in tumor cells promotes a highly plastic phenotype that facilitates invasion and colonization of the peritoneum, among others by generating a pro-tumoral immune microenvironment (**Figure 6F**).

## Discussion

The peritoneum is the second most common metastatic site in PDAC, with peritoneal metastasis contributing significantly to the devastating prognosis of patients. Our work identifies the extracellular protein HAPLN1 as a driver of tumor cell plasticity, promoting EMT, stemness, invasion and immunosuppression in a paracrine manner.

Cellular plasticity, as a key feature of tumor progression, metastasis and therapy resistance, is part of the latest list of “cancer hallmarks” (4). ECM composition and remodeling are important factors influencing cellular plasticity. Here we addressed the function of the ECM modifier HAPLN1, a linker protein for HA and proteoglycans. The reasons to investigate HAPLN1 functions were based on the finding that HAPLN1 was one of the most upregulated HA-related genes in PDAC patients and on the fact that HA is a critical component of the tumor microenvironment, modifying EMT and invasion capacity of cancer cells. Our study showed an impact of HAPLN1 on various different cell types *in vitro* and *in vivo*. In KPC cells, HAPLN1 expression resulted in the upregulation of EMP, stemness, invasion and peritoneal colonization. On top of the upregulation of several mesenchymal transcription factors like *Twist1* or *Snai2*, we found PDPN upregulated in KPC-HAPLN1 cells. PDPN is a coreceptor of the HA receptor CD44 and has been shown to induce directional, ameboidal cell migration (37).

Further, our data indicate that tumor cells with high HAPLN1 expression become more independent of CAFs as matrix producers. These cells are more adapted to survive in suspension in the peritoneal cavity, after detachment from the tumor and its stroma. Interestingly, it was already shown that also circulating tumor cells (CTCs) in pancreatic cancer expressed high levels of ECM proteins compared to their counterparts in the solid tumor, which might protect them from immune cells, similar to platelets, and ease their survival by the ability to form clusters (38).

Regarding the impact on the immune system, CAFs in HAPLN1-rich environment shifted from ECM producers in KPC tumors to more immunomodulatory, which combined with the altered transcriptome of tumor cells could be the cause for the changes in the immune compartment within the peritoneal cavity. Both, CAFs and tumor cells have been shown to act as immune modulators in cancer, promoting tumor progression by creating favorable niches (39,40). For instance, by the expression of CD47, a “don’t eat me” signal, tumor cells evade phagocytosis by macrophages (41). Interestingly, we found that *Cd47* levels significantly increased from KPC to KPC-HAPLN1 and even further in KPC-HAPLN1 suspended in the PL. We also found several modulators of inflammation upregulated in KPC-HAPLN1 cells, e.g. *Ccl2, Lcn2* or *Cxcl5*, which have been shown to regulate immune cell infiltration, contributing to disease progression (42–44).

In PDAC, several immune cell types have been described to promote different stages of tumor progression, including B cells (45), TAMs (46) and others. Data of immune cells in the peritoneal cavity during PDAC progression is however sparse and inconclusive. In other tumor entities colonizing the peritoneum, macrophages were found to be a key promoter of dissemination into the peritoneum and survival of detached cells (13,47). In KPC-HAPLN1 tumor-bearing mice the presence of macrophages was significantly increased, with a more alternative, antiinflammatory expression profile. This could form the base of an immune suppressive environment that would enable the immune evasion of disseminated KPC-HAPLN1 tumor cells in the peritoneal cavity, promoting tumor progression. The impact of the other immune cell types that were differentially recruited could not be unraveled in detail and needs further investigation.

Assessing HAPLN1 levels in PDAC patients could have diagnostic significance. It could serve to identify patients with a predisposition for peritoneal metastasis and open the possibility for early changes in treatment strategies. Since most patients that acquire peritoneal metastasis throughout disease progression acquire it after primary tumor detection (9 % of patients with peritoneal metastasis at time of detection vs. 25-50 % at time of death (3)) the classification by HAPLN1 could reduce the acquisition of peritoneal metastasis, thereby improving patients’ outcome. Nevertheless, more investigation is needed to gain better mechanistic and overall understanding of HAPLN1 action within the tumor microenvironment.

Taken together, we could demonstrate that HAPLN1 expression in tumor cells promotes a highly plastic phenotype that facilitates invasion and colonization of the peritoneum, among others by creation of a pro-tumoral immune microenvironment.

## Methods

### qRT-PCR

For transcriptomic analysis, cell lines were lysed, and RNA was isolated according to manufacturer’s protocol of the innuPREP RNA Mini Kit (Fisher Scientific, 10489573). Bulk tumor samples were homogenized with metal beads and RNA isolated with help of Trizol reagent (Life Technologies, 15596026) and isopropanol precipitation. RNA of sorted cells was isolated using the PicoPure RNA Isolation Kit (Life Technologies, KIT0214). Reverse transcription was performed with the High-Capacity cDNA Reverse Transcription Kit (Life Technologies, 4368814) or the SuperScript IV VILO Master Mix (Life Technologies, 11766050). qRT-PCR was executed using Power SYBR Green PCR Master Mix (Life Technologies, 4368708) and the primers listed in **Table 1**. Relative expression levels were calculated with the 2^-ΔΔCt^ method after normalization to the house keeping genes *Cph* or *Hprt*.

**Table 1:**
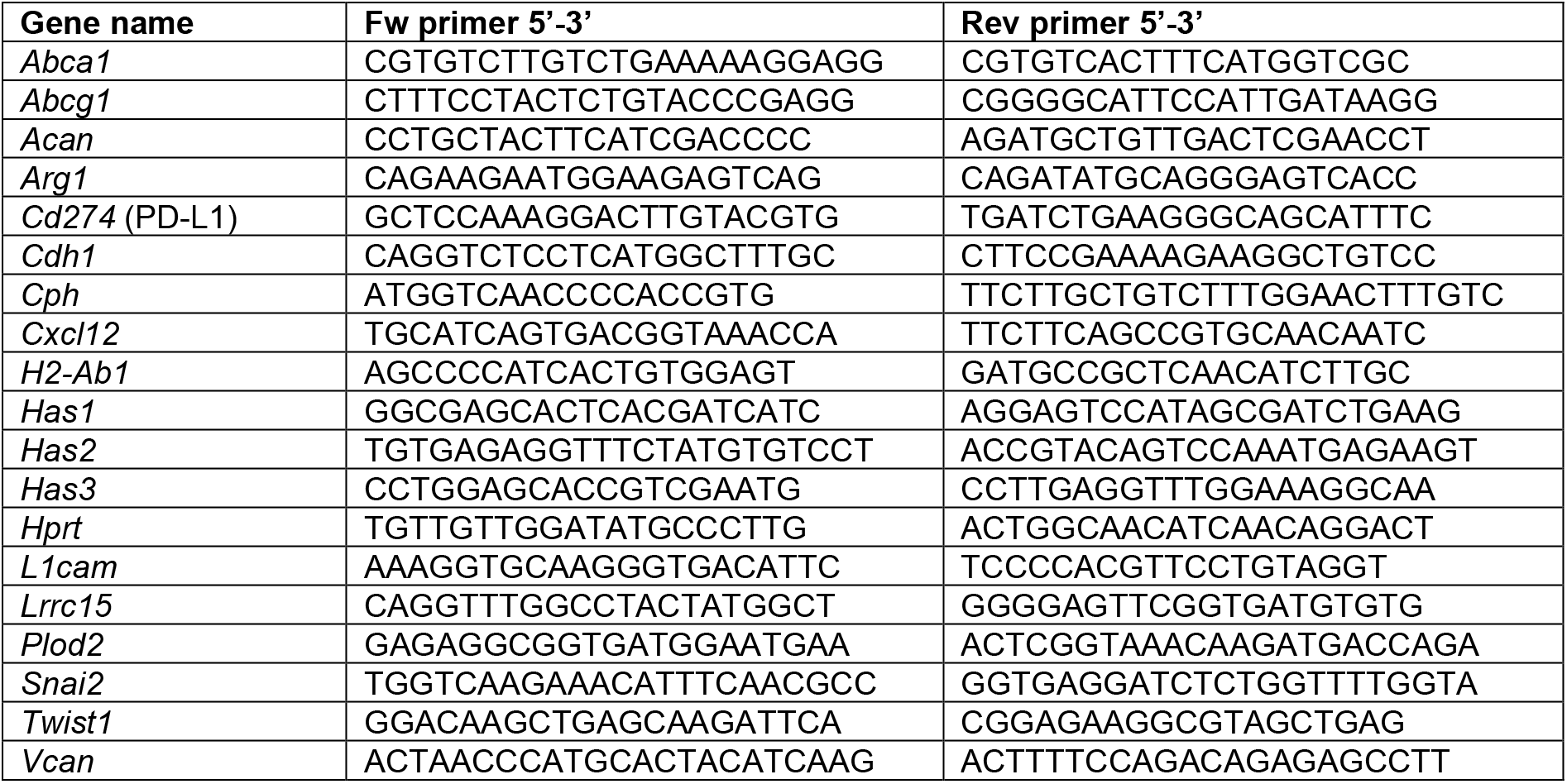
Primers for qRT-PCR

### Western blot

For Western blot analysis, cells were lysed with the Cell Lysis Buffer (Cell Signaling, 9803S). Subsequentially, samples were run on a SDS-PAGE and blotted onto PVDF membranes (Merck Chemicals, ISEQ00010) overnight. After blocking, primary antibodies were incubated overnight in 5 % milk-TBST using the following concentrations: anti-HAS2 (1:500, Abcam, ab140671), anti-VCP (1:5000, Abcam, ab109240), anti-ACTB (1:5000, Sigma Aldrich, A5441), anti-LRRC15 (1:1000, Abcam, ab150376), anti-E-Cadherin (1:2000, BD Biosciences, 610181), anti-HAPLN1 (1:1000, Bio-Techne, AF2608). Secondary antibodies goat anti-rabbit HRP, rabbit anti-mouse HRP (1:2500, DAKO, P0448, P0260) or donkey anti-goat HRP (1:2500, R&D, HAF109) were incubated for 1 h prior to detection with SuperSignal West Pico Plus Chemisubstrate (VWR International, PIER34577).

### Cell culture maintenance

All cells used for *in vitro* culture were cultured in DMEM, low glucose (high glucose for HEK293A+T cells), GlutaMAX Supplement (Life Technologies, 21885108) supplemented with 10 % FCS (VWR International, S181B) and 1 % Pen/Strep (Life Technologies, 15140122). For 2D experiments, cells were starved for 48 h prior to analysis.

### Mouse experiments

All mouse experiments were approved by the local authorities (RP Karlsruhe and DKFZ) and carried out following their legal requirements. For mouse experiments, female C57BL/6J mice were ordered from Janvier at 9-11 weeks of age. 1 million KPC or KPC-HAPLN1 cells expressing Luciferase and RFP were injected intraperitoneally into mice. Tumors grew for 11 days. Peritoneal lavage was performed to remove cells floating in the peritoneum by injection and subsequent removal of PBS into the peritoneal cavity after euthanization. For flow cytometry and cell sorting, solid tumors were digested using 2,5 mg/ml Collagenase D (Sigma Aldrich, 11088866001), 0,5 mg/ml Liberase DL (Sigma Aldrich, 5466202001) and 0,2 mg/ml DNase (Sigma Aldrich, 10104159001) in 10 % FCS DMEM for 45 min at 37 °C after mincing thoroughly. If peritoneal lavage contained erythrocytes, red blood cell lysis was performed with RBC Lysis Buffer (Life Technologies, 00-4333-57).

### Flow cytometry and Cell Sorting

Analysis by flow cytometry was performed on a BD LSR Fortessa, while BD FACS Aria/BD FACS Aria Fusion Sorters were used for cell sorting. The listed antibodies or dyes in **Table 2** were incubated for 30 mins with the cells before washing and analysis/sorting. For intracellular staining, cells were first stained with extracellular staining, then fixed, permeabilized and intracellular staining for 20min was performed. Compensation was carried out with UltraComp eBeads Compensation Beads (Life Technologies, 1-2222-42). FlowJo Software was used for analysis of the acquired samples.

**Table 2:** Flow cytometry and cell sorting antibodies

| Target with Conjugate | Dilution | Company | Cat. No. |
| --- | --- | --- | --- |
| B220-APC | 1:100 | BioLegend | 103211 |
| CD11b-BUV805 | 1:800 | BD Biosciences | 741934 |
| CD11b-PE-Cy7 | 1:800 | BD Biosciences | 552850 |
| CD19-Alexa700 | 1:50 | BD Biosciences | 557958 |
| CD25-PE | 1:200 | Life Technologies | 12-0251-82 |
| CD31-APC | 1:100 | BD Biosciences | 561814 |
| CD3-PerCp-Cy5.5 | 1:50 | BioLegend | 100217 |
| CD45-PerCp-Cy5.5 | 1:200 | BD Biosciences | 550994 |
| CD45-FITC | 1:400 | BD Biosciences | 553079 |
| CD4-PE-Cy7 | 1:300 | Life Technologies | 25-0041-82 |
| CD8-APC-Cy7 | 1:100 | BD Biosciences | 557654 |
| DAPI (stock: 10,5 mM) | 1:10.000 | Life Technologies | D3571 |
| F4/80-Alexa Fluor 488 | 1:100 | Life Technologies | 53-4801-82 |
| FoxP3-Alexa Fluor 488 | 1:50 | BD Biosciences | 560403 |
| Ly6C-PE | 1:200 | BD Biosciences | 560592 |
| Ly6G-APC-Cy7 | 1:100 | BD Biosciences | 560600 |
| MHC-II-BV510 | 1:100 | BD Biosciences | 742893 |
| PDPN-APC-Cy7 | 1:100 | BioLegend | 127417 |
| SiglecF-BUV395 | 1:100 | BD Biosciences | 740280 |

### Generation of (stable) cell lines

Stable overexpression was achieved by lentiviral infection of KPC cells. For HAPLN1 overexpression the sequence coding for HAPLN1 was inserted into a pLenti6-V5dest plasmid. The plasmid pLL3.7 (Addgene #11795) was used for eGFP, while EF.CMV.RFP plasmid (Addgene #17619) or pLenti-RFP-Puro (Hölzel, LTV-403) was used for RFP labeling. The plasmid of interest was transfected together with the plasmids for packaging (psPAX2) and envelope (pMD2, encoding for VSV-G) into HEK293T cells. Cells were infected and successful integration into the genome was guaranteed by selection with Blasticidin-Hydrochloride (Sigma Aldrich, 15205) for a minimum of 10 days. For control cells, an empty pLenti6-V5dest plasmid was used. eGFP and RFP positive cells were sorted out to receive a homogeneously marked population.

For adenoviral infection, plasmids encoding for eGFP and mCherry were amplified in HEK293A cells and cells of interest were infected afterwards. Analysis was carried out no later than 96h post infection to prevent loss of overexpression.

### Spheroid formation and invasion assay

For 3D culture, 5.000 cells were seeded in a 96 well ultra-low attachment plate (NEO Lab, 174929) in DMEM + 10 % FCS. After 96 h, pictures of the formed spheroids were taken with a Zeiss Cell Observer. For analysis by mRNA or flow cytometer, cells were seeded in 6 well plates coated with Poly(2-hydroxyethyl methacrylate) (Poly-HEMA, Sigma Aldrich, P3932) and afterwards lysed, or digested with 0,05 % Trypsin (Life Technologies, 25300054) for 5 min respectively. Subsequential to digestion, cells were stained with DAPI for analysis by flow cytometry. For invasion assay, spheroids were formed for 48 h in 96 well plates before treating them for 24 h with different inhibitors (HASi (4-Methylumbelliferrone (4-MU), 500 μM, Sigma Aldrich, M1381), FAKi (GSK2256098, 10 μM, Cayman, Cay22955-1), STAT3i (Stattic, 25 μM, Abcam, ab120952), PI3Ki (LY294002, 10 μM, BioTrend, HY-10108), MEKi (PD98059, 50 μM, New England Biolabs, 9900L) or DMSO as control. Then, half of the medium was replaced with Matrigel (Fisher Scientific, 11523550) and the different inhibitors/DMSO before polymerization to keep the concentration as indicated. For cocultures, equal amounts of KPC and KPC-HAPLN1 cells expressing either RFP/mCherry or GFP were seeded. 48 – 120 h after addition of Matrigel, pictures of invasion were taken by a Zeiss Cell Observer or a Zeiss LSM710 ConfoCor3.

### BMDM extraction

Bone marrow progenitors were isolated from the leg bones (tibia and femur) of wildtype mice by flushing the bones with PBS. Cells were then cultured on untreated plates for 7 days adding 10 ng/ml recombinant murine M-CSF (Peprotech, AF-315-02-100) at day of isolation. Afterwards, cells were counted and seeded for experiments.

### Immunofluorescence

For staining of formalin-fixed paraffin-embedded tumors were deparaffinated. Antigen retrieval in pH=9 was used for pSTAT3 and CD31 stainings, pH=6 for CD45 staining and treatment with Proteinase K for PDPN staining. Afterwards, slides were blocked, and then incubated ON with anti-CD31 (1:50, Abcam, ab28364), anti-pSTAT3 (1:200, Cell Signaling, #9145), anti-CD45 (1:50, BD Bioscience, 553076) or anti-PDPN (1:400, Abcam, ab11936). After washing, slides were incubated for 2 h at RT with anti-αSMA antibody (1:200, Sigma Aldrich, C6198-100UL) and Goat anti Rabbit Alexa Fluor 647 (1:200, Life Technologies, A21245), Goat anti Rat Alexa Fluor 647 (1:200, Life Technologies, A21247), or Goat Anti-Syrian hamster IgG H&L Alexa 647 (1:200, Abcam, ab180117). After washing, nuclei were stained using DAPI at 2,1 nM. Pictures were taken with the Zeiss AxioScan.Z1. Quantification was performed using Fiji software.

### Scratch Assay

For the scratch assay, 100.000 cells were seeded into the wells of a cell view cell culture slide (Greiner Bio-One, 543078). The next day, cell were starved and a small pipette tip was used to generate a scratch through the well. Immediately after, cells were placed into a Zeiss Cell Observer with incubation chamber, keeping the conditions at 5 % CO2 and 37 °C. Pictures were taken every 30min for a total time span of 15 hours. Migration of the cells was analyzed with the Fiji Software.

### scioCyto Cytokine array of serum

Blood was taken from the *vena facialis* 11 days after tumor injection and collected in a microvette LH (Sarstedt, 201.345) to receive the serum. Sciomics GmbH running the scioCyto Cytokine array performed the analysis of the serum of tumor bearing and control mice.

### *ex vivo* luminescence

For *ex vivo* luminescence was measured on cells of the peritoneal lavage using Luciferase Assay System (Promega, E1500) and a CLARIOStar plate reader.

### RNA sequencing

Libraries and multiplexes were prepared with NEBNext Multiplex Oligos for Illumina (New England Biolabs, E7600S) and NEBNext Single Cell/Low Input RNA Library Prep Kit for Illumina (New England Biolabs, E6420S). NextSeq 550 PE 75 HO (Illumina) RNA sequencing was performed by the Genomics & Proteomics Core Facility of the German Cancer Research Center (DKFZ, Germany).

### Gene Set Enrichment Analysis (GSEA)

To investigate differentially expressed genes, Gene Set Enrichment Analysis (GSEA, (49,50)) was performed. p values were assessed using 1000 permutations for each gene set and corrected with the false discovery rate (FDR) method. When analyzing microarray data, the mean of all probes for the same gene was used to divide patients into high/low cohorts. The following publicly available data sets of PDAC patients were used for our analyses: NCBI-GEO GSE62452 and the data from Cao et al. 2021 (22,23).

### Bioinformatic analysis of RNA-sequencing data

Bioinformatics analysis of RNA-sequencing data sets was performed using the software R using the DESeq2 package. Plots were created using ggplot2 or EnhancedVolcano. apeglm package was used for LFC shrinkage.

### Statistics

Graphs represent mean ± SD with each data point depicting a biological replicate. Statistical analysis was performed using the GraphPad Prism 9 software. For *in vitro* experiments, Gaussian distribution was assumed, and two-tailed parametric unpaired t-test was applied, while the non-parametric Mann-Whitney test was used for *in vivo* and *ex vivo* experiments. Data was considered as statistically significant if p<0.05. For high throughput analysis (RNAseq, scioCyto Cytokine array) p_adj_ was calculated and results were considered significant if p_adj_<0.05.

### Schemes

Schemes were generated using BioRender.com.

## Acknowledgements

We would like to thank Stephen Konieczny, from Purdue University, Indiana, for providing us the KPC cell line. Additionally, we thank the Light Microscopy core facility, the High Throughput Sequencing Unit of the Genomics and Proteomics Core Facility and the Omics IT and Data Management (ODCF), the Flow Cytometry core facility and animal caretakers of the German Cancer Research Center (DKFZ) for providing excellent services. We would like to thank Damir Krunic (DKFZ, Light Microscopy Core Facility) for his help with FIJI software data analysis.

This work was funded by the Deutsche Forschungsgemeinschaft (DFG) project 394046768 -SFB1366 projects C4 (to A.F.), DFG project 419966437, Deutsche Krebshilfe project 70113888, MCIN/AEI/ 10.13039/501100011033 (PID2020-117946GB-I00 and RYC2019-027937-I) and RYC2019-027937-I) (to J.R-V.). The Science Ministry of Spain or the Health Ministry (ISCIII) receives support from the EU and its ERDF program. Part of the equipment used in this work has been funded by Generalitat Valenciana and co-financed with ERDF funds (OP ERDF of Comunitat Valenciana 2014-2020).

## Author Contributions

Conception and design: L.W., F.D-R., A.F., J.R-V.; Acquisition of data: L.W., F.D-R., N.V-S., E.D., I.M., M.V., J.G.; Analysis and interpretation of data: L.W., F.D-R., N.V-S., E.D., E.E., E.A-S., R.M., A.T., A.F., J.R-V.; Writing and revision of the manuscript: L.W., F.D-R., J.R-V., A.F.; Study supervision: J.R-V., A.F.

