## Supplementary figures and images for "HAPLN1 is a driver for peritoneal carcinomatosis in pancreatic cancer"

### Suppl.Video1.gif

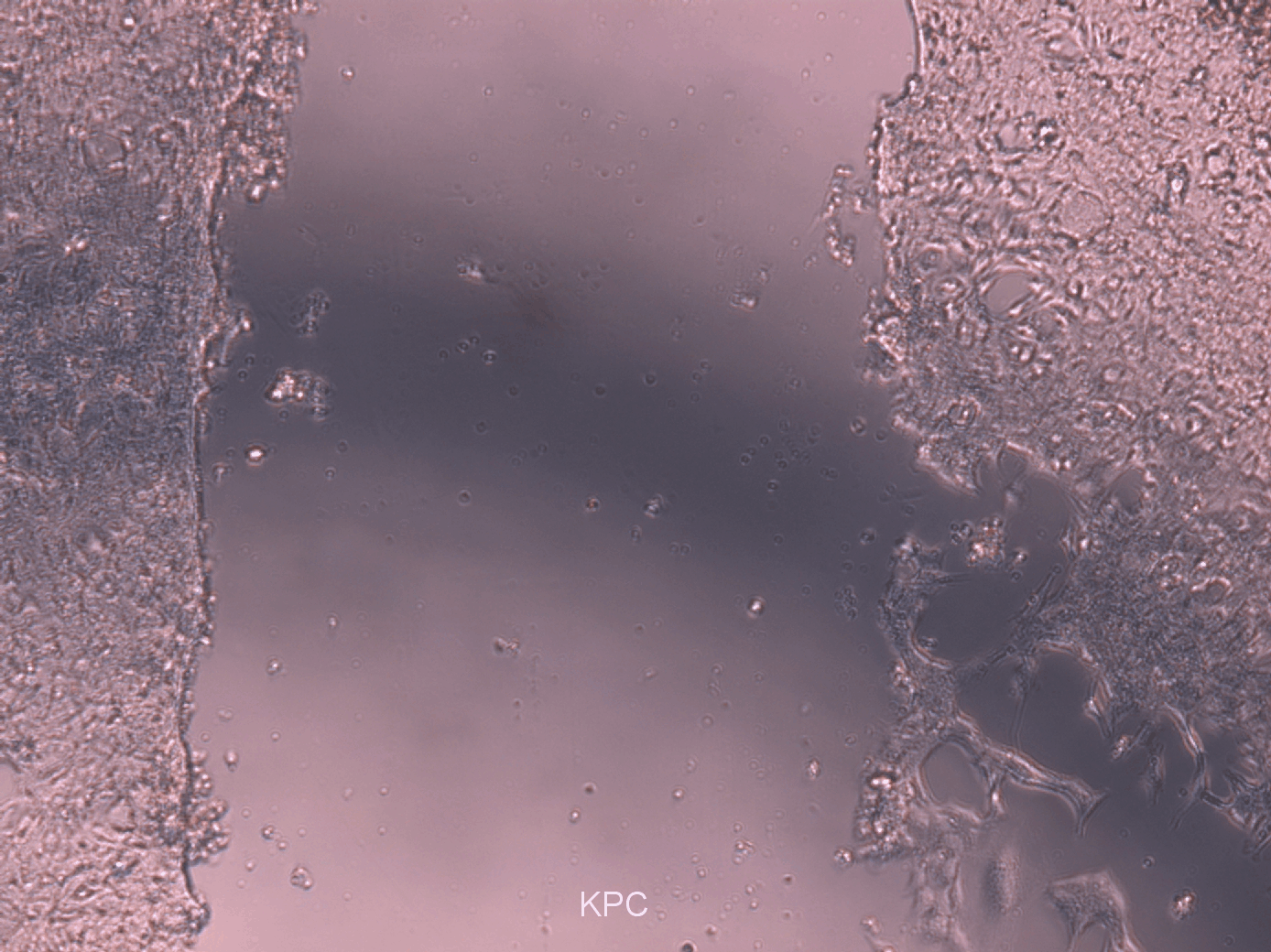

### Suppl.Video2.gif

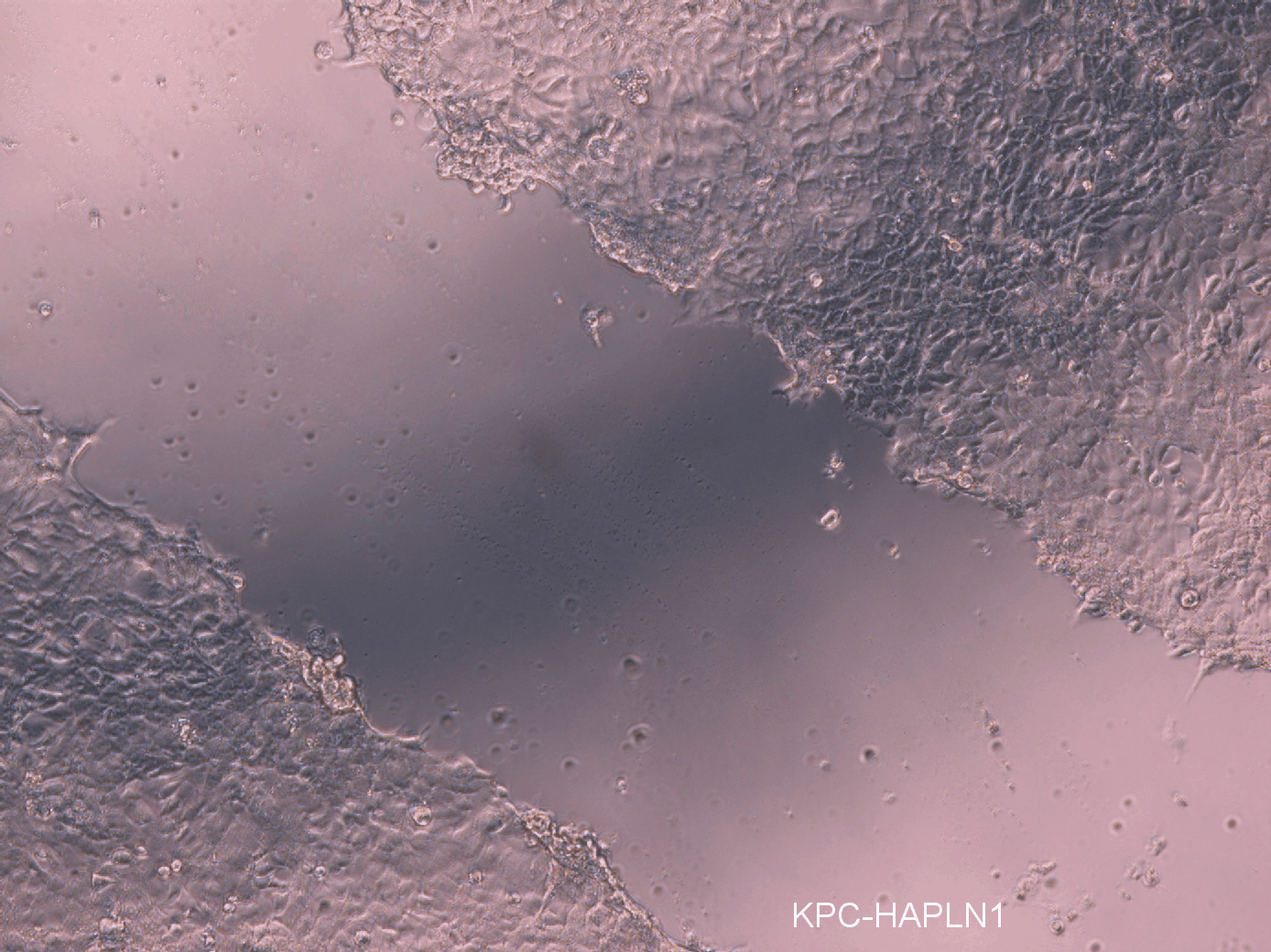
