## Supplementary material for "HAPLN1 is a driver for peritoneal carcinomatosis in pancreatic cancer": Suppl. Figure

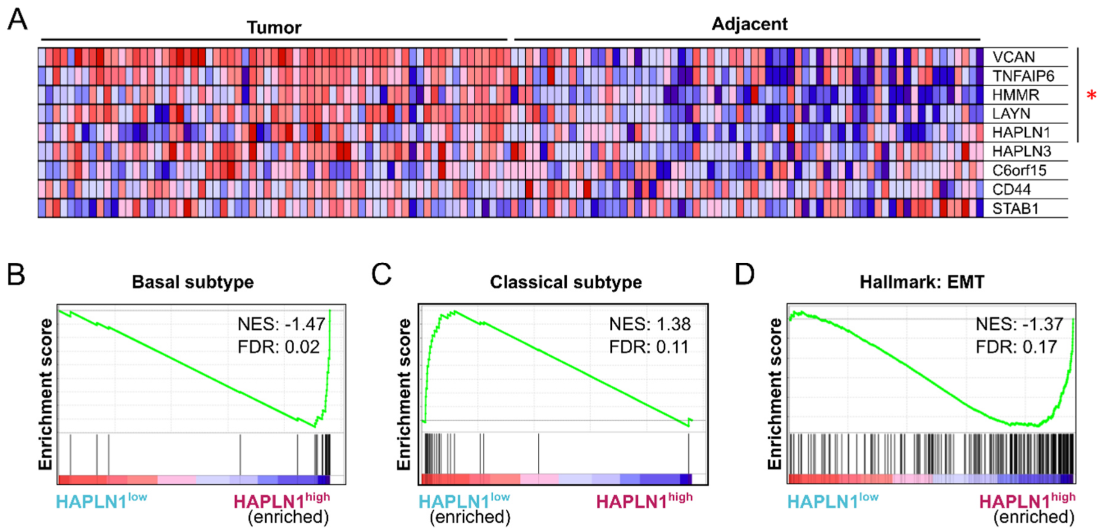


**Supplementary Figure 1: HAPLN1 is upregulated in PDAC tissue and associated with worse disease outcome**

**A**. GSEA on GSE62452 (n=61 normal adjacent and n=69 PDAC tumor samples, (23)) on "Gene ontology for Hyaluronic Acid binding". Leading edge genes significantly deregulated are marked with red star. **B-D.** GSEA on GSE62452. Before analysis PDAC patients were divided in *HAPLN1* high/low according to mean *HAPLN1* expression. Gene sets of "Basal subtype" (**B**), "Classical subtype" (**C**) or "Hallmark of Epithelial-to-Mesenchymal transition" (**D**). n=69.

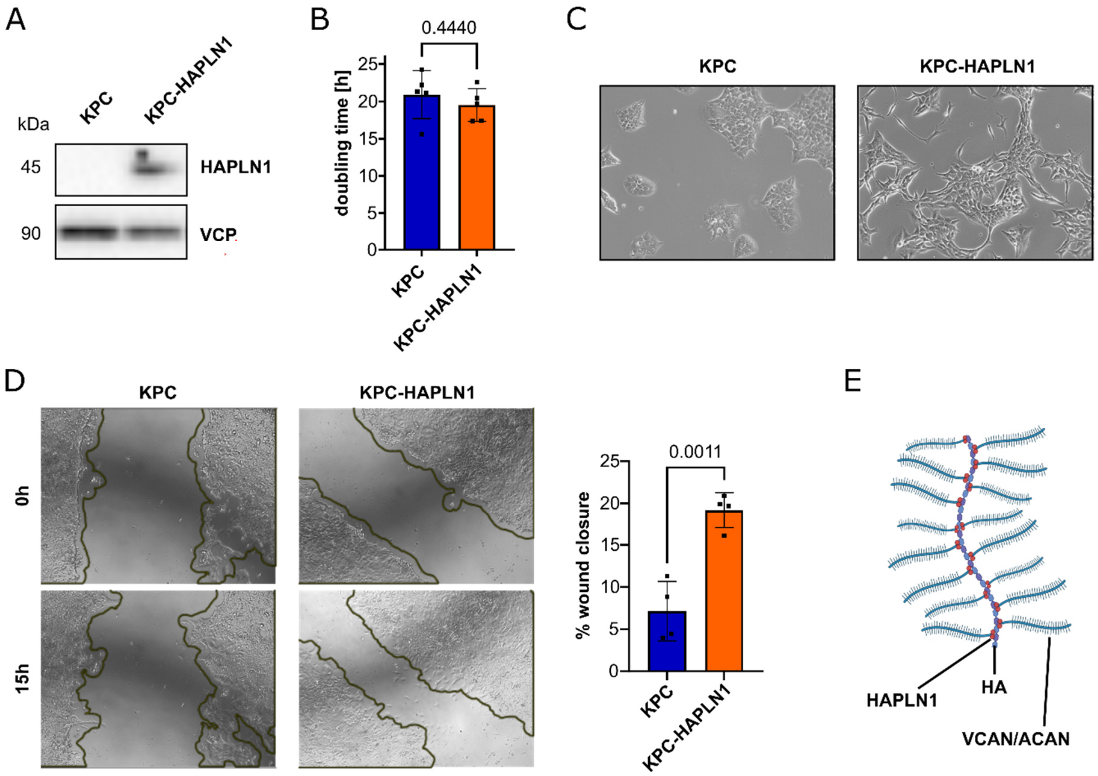


**Supplementary Figure 2: HAPLN1 induces EMT and ECM remodeling *in vitro***

**A**. Confirmation of HAPLN1 overexpression by Western blot analysis with VCP as house keeper. Representative image shown. **B**. Doubling time of KPC and KPC-HAPLN1 assessed by cell counting. n=4. **C**. Cellular morphology of stably transfected cells. Scale bar: 50 µm. **D**. Scratch assay of KPC and KPC-HAPLN1 cells for 15 h. Closure was monitored by live cell imaging. Borders in the beginning (0 h) and in the end (15 h) are marked. n=4. **E**. Schematic overview of HAPLN1 function as crosslinker of HA and proteoglycans Aggrecan (ACAN) or Versican (VCAN). Graphs display mean±SD. Data points mark independent biological replicates. For all data, unpaired two-tailed T test was applied.

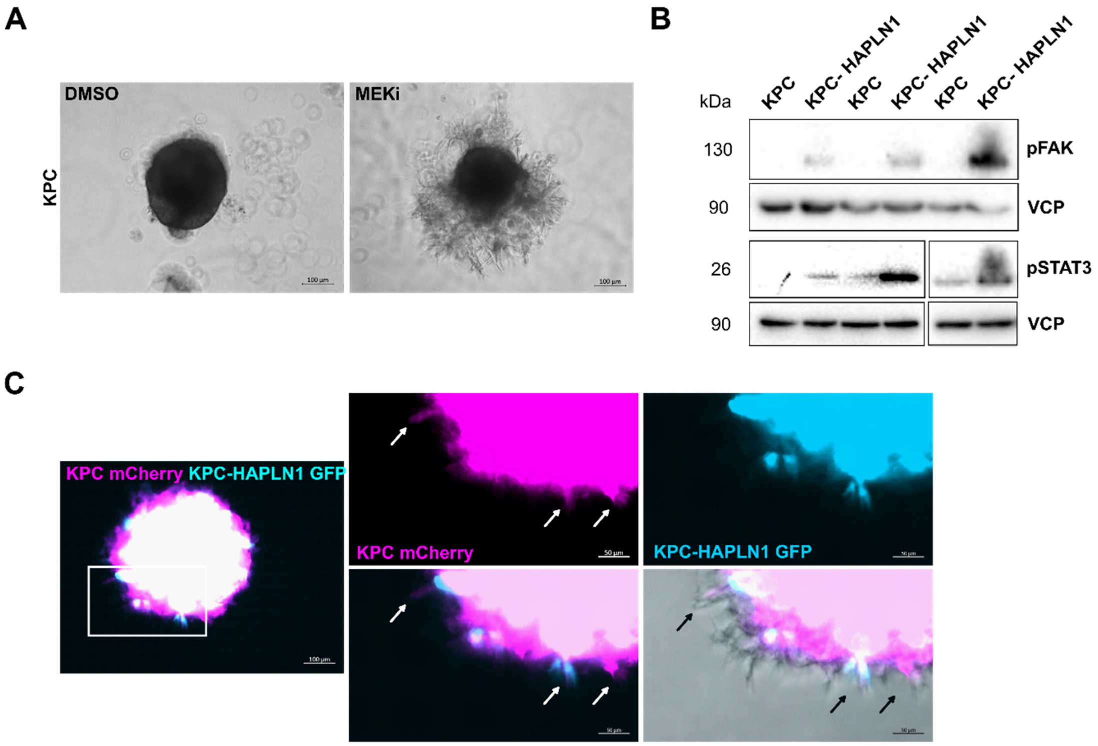


**Supplementary Figure 3: HAPLN1 fuels invasion and acts in a paracrine manner**

**A**. KPC spheroids embedded in Matrigel after 48 h. Spheroids were pre-treated with DMSO or MEK inhibitor (MEKi; 50 µM PD98059). Representative image shown. Scale bar: 100 µm. **B**. Western blot analysis of STAT3 and FAK phosphorylation of untreated KPC or KPC-HAPLN1 cells. n=3. **C**. KPC and KPC-HAPLN1 cells were labeled by adenoviral infection with mCherry or GFP. Invasion was assessed after 48 h. Representative image shown. Scale bar: 100 µm, zoom: 50 µm.

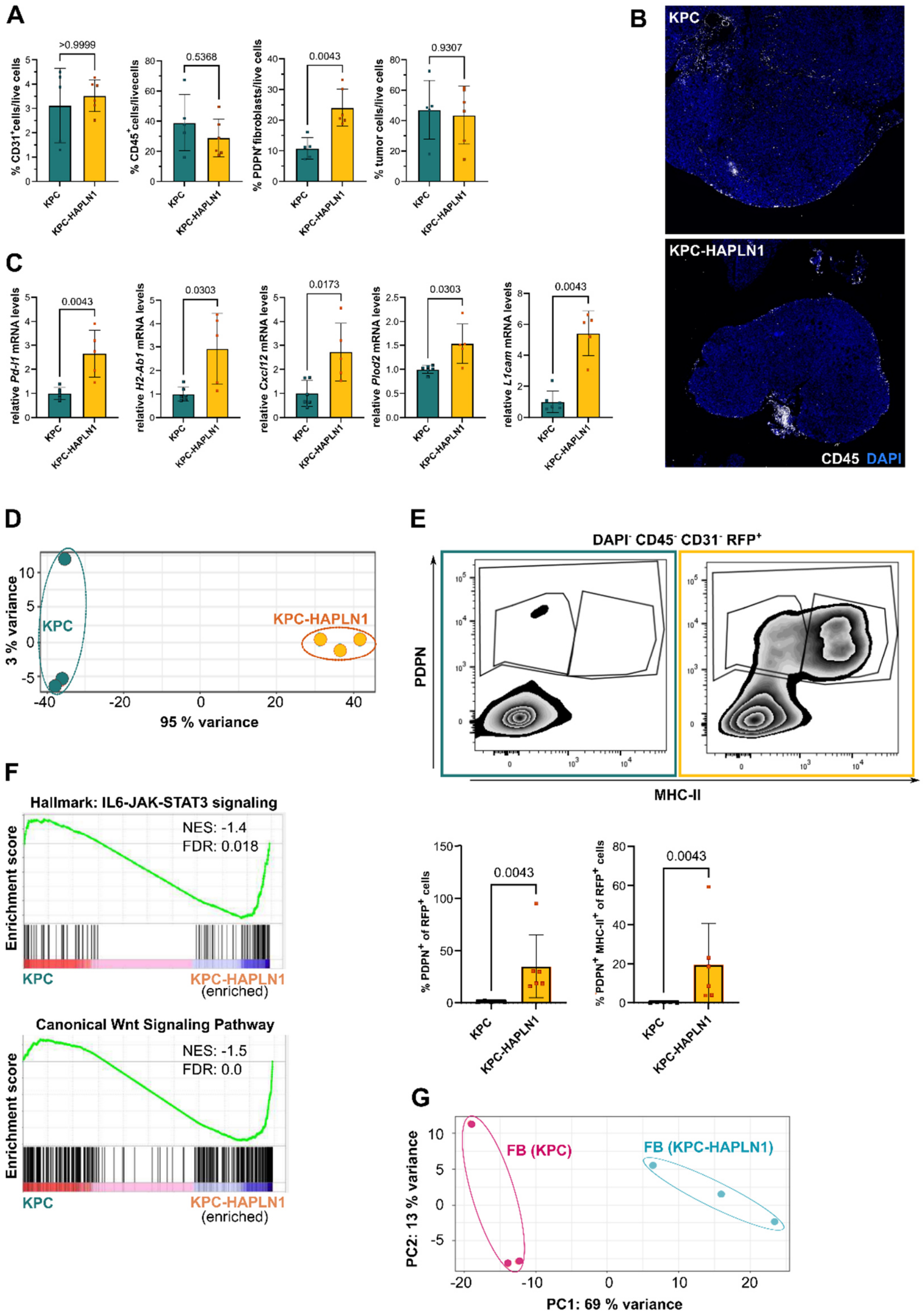


**Supplementary Figure 4: HAPLN1 induces tumor cell hyperplasticity *in vivo***

**A**. Analysis of tumor composition after digestion by flow cytometry. n=5-6. **B**. Immunofluorescence staining for CD45. One tumor nodule shown as representative image. **C**. qRT-PCR analysis of whole tumor mRNA. n=5-6. **D**. Principal component analysis (PCA) of isolated tumor cells send for RNA sequencing. Replicates of the same group are marked with same color. PC=principal component. **E**. Flow cytometric analysis of digested tumor samples. Representative plots of MHC-II and PDPN expression by tumor cells and quantification. n=5-6. **F**. GSEA of RNAseq data of isolated tumor cells on "Hallmark of IL6-JAK-STAT3 signaling" and "Canonical Wnt Signaling Pathway" gene sets. **G.** Principal component analysis (PCA) of isolated fibroblasts (FB) of KPC or KPC-HAPLN1 tumors. Data points represent independent biological replicates, data shown is mean±SD. Non-parametric Mann-Whitney U test was applied.

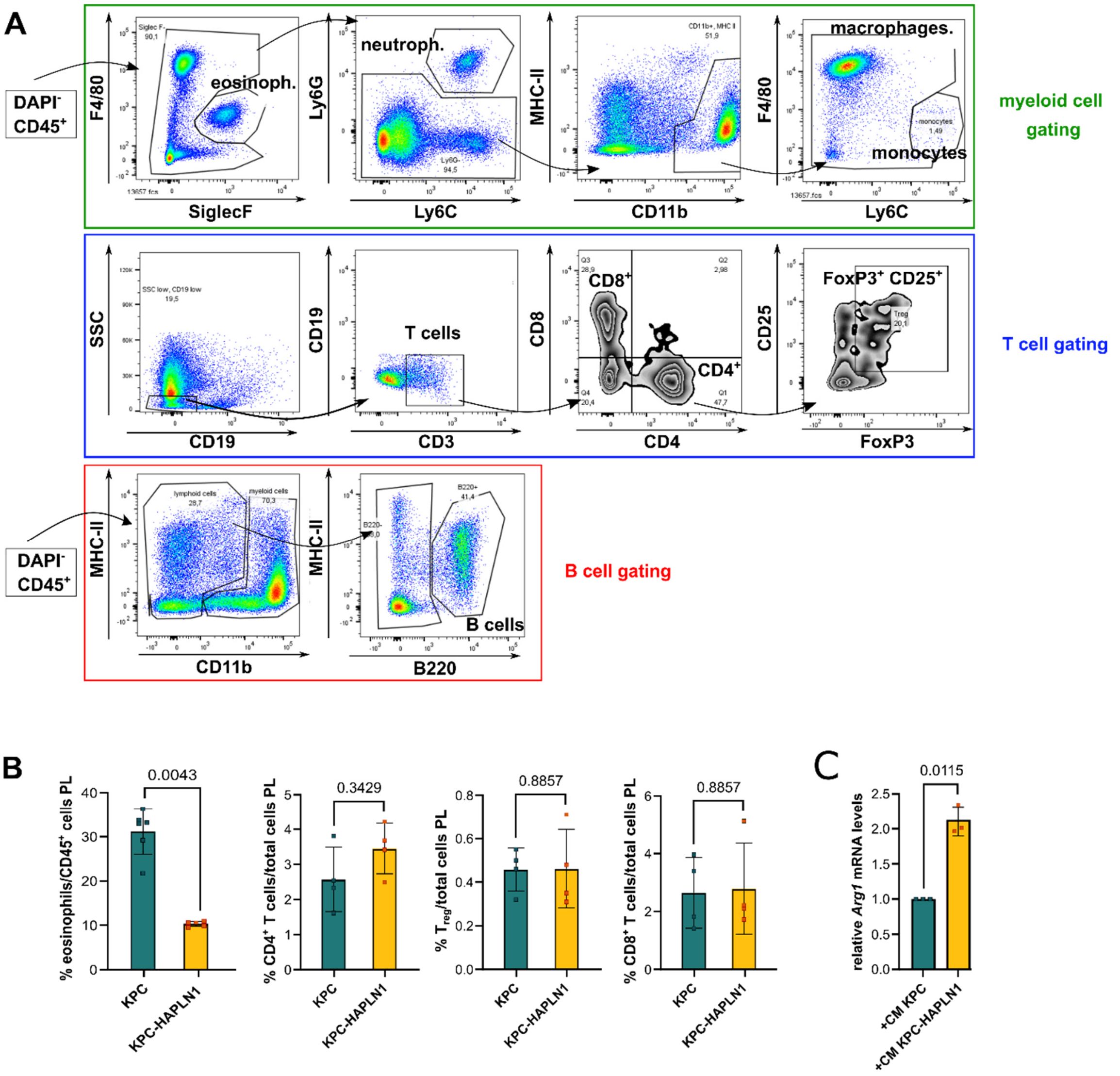


**Supplementary Figure 5: HAPLN1 modifies the immune microenvironment**

**A.** Gating of flow cytometric analysis of immune cell populations. Representative sample shown. **B.** Immune cell populations in the peritoneum assessed by flow cytometry. n=5-6 for eosinophils, n=4 for T cells. **C.** Bone marrow-derived macrophages (BMDMs) were treated with conditioned medium (CM) of KPC or KPC-HAPLN1 tumor cells for 24 h. Gene expression of *Arg1* was analyzed by qRT-PCR. n=3. Data points represent independent biological replicates, data shown is mean±SD. Non-parametric Mann-Whitney U test was applied for panel B, paired two-tailed T test for panel C.

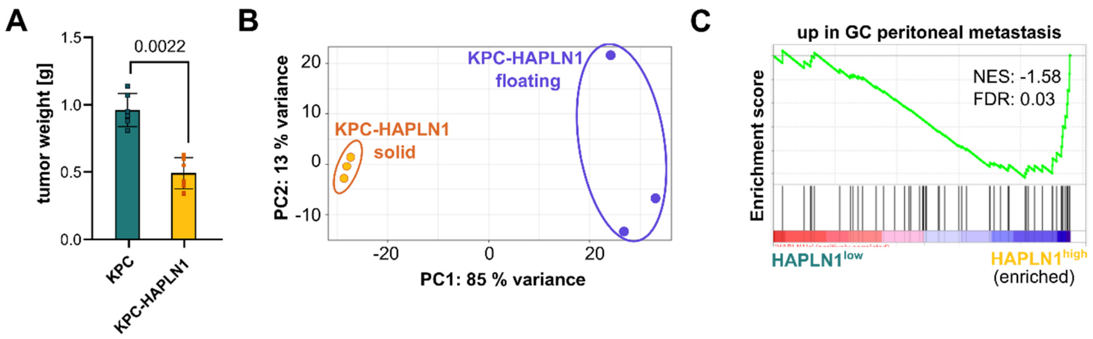


**Supplementary Figure 6: HAPLN1 facilitates peritoneal colonization by tumor cells**

**A.** Solid tumor weight after 11 days of tumor growth. n=6. **B.** KPC-HAPLN1 cells from solid tumor and in suspension were isolated and send for RNAseq. Principal Component analysis (PCA) of the samples. n=3. **C.** GSEA of GSE62452, dividing patients into *HAPLN1* high/low according to their mean *HAPLN1* expression. A gene set for genes upregulated in gastric cancer peritoneal metastasis was analyzed (36). n=69. Data points represent independent biological replicates, data shown is mean±SD. Non-parametric Mann-Whitney U test was applied for panel A.
